# A Miniaturized 3D-Printed Pressure Regulator (*μ*PR) for Microfluidic Cell Culture Applications

**DOI:** 10.1101/2022.04.03.486540

**Authors:** Meng-Chun Hsu, Mehran Mansouri, Nuzhet N.N. Ahamed, Indranil M. Joshi, Adeel Ahmed, David A. Borkholder, Vinay V. Abhyankar

## Abstract

Controlled fluid flows are the hallmark feature of microfluidic culture systems and provide precise definition over the biophysical and biochemical microenvironment. Flow control is commonly achieved using displacement-based (e.g., syringe or peristaltic pumps) or pressure-based techniques. These methods offer complex flow capabilities but can be challenging to integrate into incubators or other confined environments due to their large form factors and accompanying peripheral equipment. Since many microfluidic cell culture studies use a single controlled flow rate to maintain or stimulate cells, a portable flow control platform that fits easily into an incubator will benefit the microfluidic community. Here, we demonstrate that a tunable, 3D printed micro pressure regulator (*μ*PR), combined with a battery-powered miniature air pump, can operate as a stand-alone pneumatic flow control platform for microfluidic applications. We detail the design and fabrication of the *μ*PR and demonstrate: i) a tunable outlet pressure range relevant for microfluidic applications (1-10 kPa), ii) highlight dynamic control in a microfluidic network, and iii) maintain human umbilical vein endothelial cells (HUVECs) in a multi-compartment membrane-based culture device under continuous flow conditions. We anticipate that our 3D-printed fabrication approach and open access designs will allow other laboratories to rapidly customize *μ*PRs to support a broad range of applications.

## Introduction

Microfluidic approaches leverage the precise manipulation of fluids to introduce unique experimental capabilities in biological applications (Wang et al. 2015; Ahmed et al. 2021; Whitesides 2006), including the defined stimulation of cultured cells (Biffi et al. 2012; Song et al. 2009), the controlled influx of chemical compounds (Gossett et al. 2012), and the introduction of secondary cell populations to the culture environment (Wartmann et al. 2015). In these systems, control over fluid flow is typically achieved via displacement-based or pneumatic pumping schemes (Zeng et al. 2015; Bong et al. 2011; Ward et al. 2005). The displacement-based mechanism is employed in syringe and peristaltic pumps. Syringe pumps use the rotary motion of mechanical screws to dispense fluid from a syringe barrel at a controlled flow rate (*Q*), while peristaltic pumps employ a cam mechanism to push or pull fluids through compliant tubing to directly control *Q* (Bong et al. 2011; Byun et al. 2014). Although syringe and peristaltic pumps are frequently used due to their robust flow control capabilities and compatibility with standardized components (e.g., syringes, fittings, and tubing), they can be challenging to integrate into confined environments. In addition, the mechanical oscillations of the rotary motor or cam mechanism can introduce undesired flow pulsations that result in cell damage (Pardo et al. 2020; Kurth et al. 2019; Kusahara et al. 2015; Wilson et al. 2016). Pneumatic pumping schemes create a defined pressure drop (Δ*P*) across microfluidic networks to drive fluid flows. The flow rate is governed by the hydraulic analogy to Ohm’s Law, *Q* = Δ*P* · *R*^−1^, where *R* is the hydrodynamic resistance of the network (Aoki et al. 2006; Dutta et al. 2006). Because of their intrinsic damping nature, pneumatic approaches are less susceptible to flow pulsations compared to displacement-based methods. However, they also require more complex peripheral equipment, such as a dedicated high-pressure air source (e.g., laboratory air), a closed-loop pressure controller, and in-line pressure/flow sensors (Wilson et al. 2016; Thurgood et al. 2018; 2019). Consequently, pneumatic methods can be difficult to integrate into cell culture environments (Mavrogiannis et al. 2016).

Both displacement and pneumatic techniques offer excellent flow control capabilities and can be programmed to dynamically adjust flow profiles, including ramped, periodic, pulsed, or even reversed flows. However, these advanced features are not often utilized in standard microfluidic culture applications where a constant flow rate is used to maintain or stimulate cultured cells (Wang et al. 2015; Thurgood et al. 2018; 2019; Marimuthu et al. 2013). The experimental need for a single controlled flow rate allows us to forgo some of the advanced flow functionalities in favor of a simple and portable pumping solution. Alternative approaches to simplify the pumping process have been widely explored. For example, a commercial palm-top refillable infusion pump was used to culture cells (iPrecio SMP101-L, Primetech, Tokyo, Japan) (Sasaki et al. 2012). However, the pump was expensive, one-time use, and could not be customized. Alternatively, passive pumping, including hydrostatic and surface tension-based methods are low-cost and easy to use, but lack long-term stability, making them unsuitable for microfluidic culturing applications (> 24 hours) (Jeong et al. 2014; Yeh et al. 2017). Microelectromechanal systems (MEMS) approaches have also been used to create microfabricated pumps (Stevenson et al. 2012; Wang et al. 2018). Although these micropumps can provide the long-term control required for lab-on-chip applications, the complexity of the fabrication procedures can make customization and implementation impractical.

To address the need for a simple but functional pumping platform, we introduce a pneumatic pumping platform that uses a 3D-printed micro pressure regulator (*μ*PR) to provide a tunable Δ*P* that controls the flow rate in a microfluidic channel network. Our *μ*PR uses a force-balance mechanism to reduce the pressure supplied by a battery-powered air pump to a controllable pressure range relevant to microfluidic applications. We detail the design and fabrication of the *μ*PR, establish dynamic pressure control and stability characteristics, and demonstrate successful culture within a membrane-based, compartmentalized microfluidic barrier model (Mansouri et al. 2022; Mccloskey et al. 2022). As 3D-printers have become broadly accessible in research laboratories (Coakley et al. 2016), we anticipate that our 3D-printed *μ*PR – with open access designs - can be fabricated and assembled in any research laboratory and tailored to achieve application-specific flow requirements.

## Materials and methods

### Fabrication of the Pressure Regulator

The structural components of the *μ*PR, including the inlet (high-pressure) and outlet (low-pressure) chambers and the pressure control component, were 3D printed using the Formlabs Form 2 stereolithography printer (Formlabs Inc., Somerville, MA, USA). Dental SG resin (Formlabs Inc., Somerville, MA, USA) was selected as the building material due to its gas-impermeable characteristics and Class I biocompatibility (EN-ISO 10993-1:2009/AC:2010). The 3D-printed parts were removed from the print platform, rinsed in 99% isopropyl alcohol, dried under pressurized air, and UV-cured for 45 minutes at 45°C in a post-processing curer (FormCure, Formlabs Inc., Somerville, MA, USA), in accordance with the manufacturer’s recommendations.

A 001 size Viton fluoroelastomer (shore 60A) O-ring (McMaster Carr, Elmhurst, IL, USA) was fitted over the connecting rod adjacent to the poppet valve of the high-pressure inlet chamber as shown in **Fig. 1a(i)**. An 8-mm-ID/10-mm-OD natural rubber (shore 70A) O-ring (McMaster Carr, Elmhurst, IL, USA) was then placed in the outer groove of the inlet chamber. The low-pressure outlet chamber, shown in **Fig. 1 a(ii)**, was placed over the inlet chamber with the connecting rod extending through the cavity to form the cross-chamber air passage. Next, a 100-*μ*m thick Kapton (Gizmo Dorks LLC, Temple City, CA, USA) was placed onto the outlet chamber as the pressure sensing diaphragm, in contact with the connecting rod. As shown in **Fig. 1a(iii)**, an O-ring was placed on top of the diaphragm to help seal the top of the outlet chamber. The pressure control component with built-in cantilever springs was then stacked onto the diaphragm. These cantilevers were 0.5 mm wide, 0.5 mm thick, and 5 mm long. An M2 nut (McMaster Carr, Elmhurst, IL, USA) was glued to the cantilever springs with epoxy adhesive (ClearWeld™ Professional, J-B Weld Company, Sulphur Springs, Texas, USA) (**Fig. 1a(iv))**. As shown in **Fig. 1(v)**, an M2 bolt was threaded into the nut. A 3D-printed pointer was added to the hexagonal socket head to create the control knob. A laser-cut, 24-position acrylic dial was attached to the pressure control component using pressure-sensitive adhesive (PSA, 3M 468MP Adhesive Transfer Tape, 3M Company, Maplewood, MN, USA). The dial provided indications for rotational positions in 15° increments. Finally, 3D-printed clamps were used to compress the outer O-rings sandwiched between the structural components and complete the assembly as shown in **Fig. 1a(vi)**. The assembled device is 12mm in diameter and 20mm in height. **Fig. 1b** shows an image of the assembled device next to a US dime for scale.

**Fig. 1.**
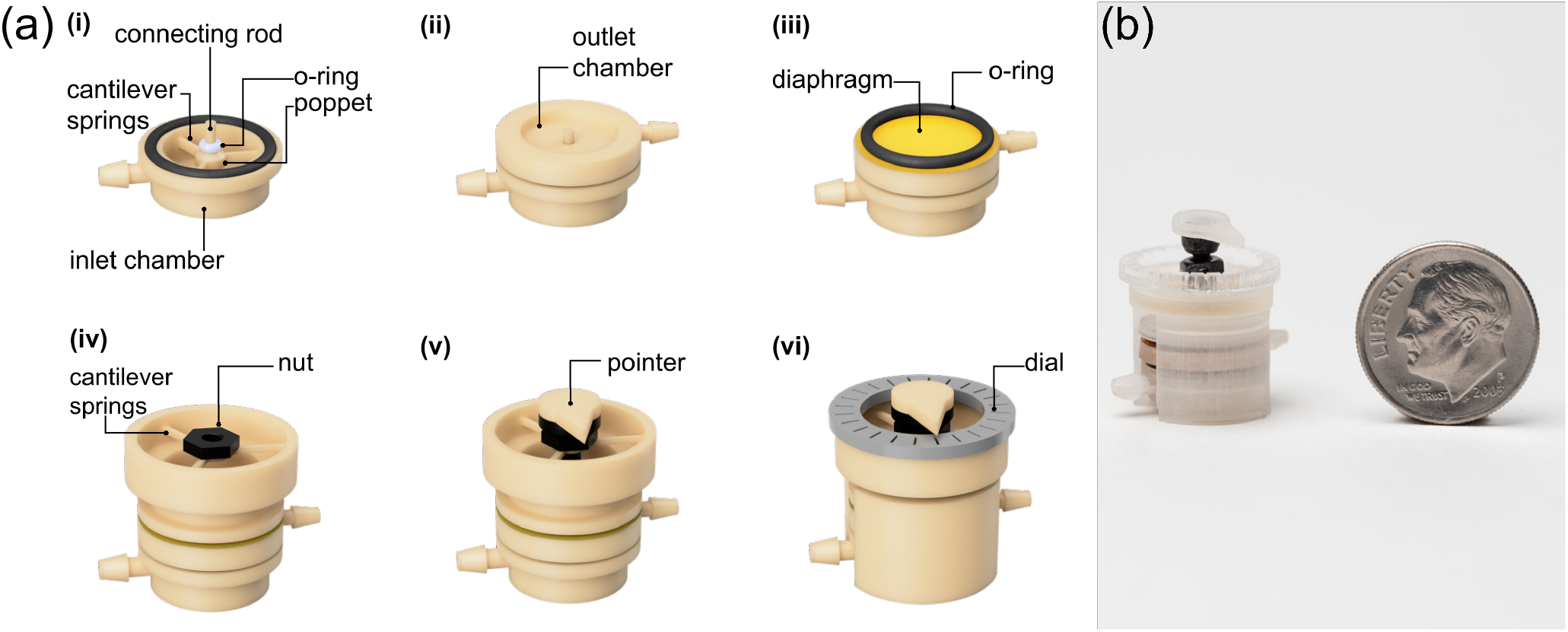
A) Schematic view of the 3D printed *μ*PR and fabrication workflow. (i) The high-pressure air inlet chamber includes a poppet valve, a sealing O-ring (white), and a connecting rod. (ii) The low-pressure air chamber is placed on top of the inlet chamber. (iii) A Kapton diaphragm (yellow) and O-ring (black) are placed atop the outlet chamber. (iv) The pressure control component, consisting of three built-in cantilevers and a threaded nut, is positioned on top of the O-ring. (v) An M2 bolt with a 3D-printed position indication pointer is threaded into the nut. (vi) The device is then sealed using two 3D-printed compression clamps to achieve an air-tight assembly (Φ12mm x 20mm) and a laser-cut position dial is added. B) Image of the assembled 3D-printed *μ*PR next to a United States dime for scale.

### Microfluidic Channel Fabrication

(Poly)dimethylsiloxane (PDMS, Sylgard 184, Dow Inc., Midland, MI, USA) microchannels were fabricated using standard soft-lithography techniques (Ahmed et al. 2021; Abhyankar et al. 2016). SU-8 2100 (Kayaku Advanced Materials, Westborough, MA, USA) was spin-coated onto a 4” silicon wafer, soft-baked, exposed to UV light through a transparency mask (CAD/Arts Services Inc., Bandon, OR, USA) to define channel features, and post-baked at 95°C. The photoresist was then developed (Kayaku Advanced Materials, Westborough, MA, USA). A rectangular PMMA ring (length = 75 mm, width = 25 mm) was attached to the wafer using PSA to create a molding cavity with a defined height. Upon attachment of the PMMA ring, the mold was then filled with degassed PDMS pre-polymer (10:1 base to catalyst ratio by mass) and cured on a hotplate for 1 hour at 80°C. The PDMS block was then removed from the mold and access ports were cored with a 1-mm biopsy punch (World Precision Instruments, Sarasota, FL, USA).

### COMSOL Flow Simulation Setup

A 3D simulation was performed using the laminar flow physics (stationary) module in COMSOL Multiphysics. Microchannel geometry (20-*μ*m height, 100-*μ*m width, and 32-cm length) was applied with the material set as water. We assigned pressures (P = 1-10 kPa) to the inlet of the microchannel geometry, while the outlet pressure was defined as atmospheric (P = 0) with suppressed backflow. The other sides of the block were assigned no slip boundary conditions.

### Pressure and Flow Rate Measurement

The general experimental setup featured a *μ*PR and a PDMS microfluidic channel (20-*μ*m height, 100-*μ*m width, and 32-cm length). We supplied pressure to the *μ*PR with a miniature DC air pump SX-2 (Binaca Pumps, Temecula, CA, USA) operating at 3V and 0.09A. The outlet of the *μ*PR was connected to a threeway connector, with one end feeding the inlet of the PDMS microfluidic channel and the other connected to a Honeywell pressure sensor (TBPDANS005PGUCV, Honeywell International Inc., Charlotte, NC, USA). Silicone tubing (2-mm ID, 5-cm length) was used to connect these components. Details of the pressure sensing setup are shown in **Fig. S1** and **Fig. S2**. The PDMS microchannel was primed with a solution of blue dye (McCormick Inc., Baltimore, MD, USA) in deionized water to improve contrast.

### Characterization of Outlet Pressure vs Control Knob Position

The aforementioned experimental setup allowed characterization of *P_out_* based on the rotational position of the control knob. The control knob was turned by 15° increments (indicated with the acrylic dial) while *P_out_* was measured. *P_out_* was then allowed to stabilize for 5-minutes for each position after turning. A full cycle of the calibration process included clockwise rotational turns (*P_out_* increased from 1 to 10 kPa) and counterclockwise turns (*P_out_* decreased from 10 to 1 kPa). 15 full cycles were used to calibrate the outlet pressure readings versus knob position. In order to quantify the stability of the regulated pressures, data was collected over a period of 1000 minutes for three designated pressures (*P_out_* = 1, 5, and 10 kPa), covering the low, medium, and high set points of the range.

### Cell Culture of HUVECs in *μ*PR Microfluidic Platform

Detailed design and fabrication of the barrier platform has been described in our previous work (Mansouri et al. 2022; Mccloskey et al. 2022). Briefly, the cell culture platform consists of the top and bottom microchannels, separated by an ultrathin nanomembrane (SiMPore Inc., Rochester, NY, USA). The nanomembrane has a thickness of 100 nm and a pore size of 60 nm. The device has a core open-well module known as the m-*μ*SiM which can be reconfigured into a fluidic device by adding a flow module into its well and sealing it magnetically using two housings with embedded magnets. The flow module was fabricated using standard soft lithography method and housings were fabricated using a laser cutter (H-series 20×12, Full Spectrum, CA, USA) (Mansouri et al., 2022). The dimensions of the top channel were, h = 200 *μ*m, w = 1.5 mm, and l = 5 mm, and the bottom channel were, h = 150 *μ*m, w = 2-6 mm, and l = 15 mm. The flow from the media reservoir connected to the inlet of the top channel using tubing and 21 gauge 21 NT dispensing tips (Jensen Global, USA).

Prior to cell seeding, the nanomembrane was coated with 5 *μ*g·cm^−2^ fibronectin (Corning Inc., Corning, NY, USA) for one hour at room temperature, and then rinsed with fresh cell media. Human umbilical vein endothelial cells (HUVECs) (Thermo Fisher Scientific, Waltham, MA, USA) were cultured in EBM-2 Basal Medium (Lonza Bioscience, Walkersville, MD, USA) supplemented with EGM-2 Endothelial Cell Growth Medium-2 BulletKit (Lonza Bioscience, Walkersville, MD, USA) and maintained in a tissue culture flask. Prior to use, cells were dissociated using TrypLE (Thermo Fisher Scientific, USA) for 3 min and centrifuged at 150 G for 5 min. After re-suspension, cells were seeded onto the membrane surface through the top microchannel and incubated for 1 hour to promote cell attachment.

The *μ*PR was set to an output pressure of 8 kPa (Δ*P* = 8 kPa), which corresponds to a media flow rate of 1 *μ*L·min^−1^ (shear stress of 0.02 dynes·cm^−2^ at cell monolayer) in the top channel for 24 hours. LIVE/DEAD stain (Thermo Fisher Scientific, Waltham, MA, USA) was used to assess cell viability based on the vendor’s protocol. Labeled cells were imaged using an Olympus IX-81 fluorescence microscope with CellSens software (Olympus, Tokyo, Japan) with constant image capture settings across the experimental sets.

### Dynamic Flow Control

A Y-shaped PDMS microchannel consisting of two 1-cm-long inlet channels and a 1-cm-long outlet channel was connected to two *μ*PRs (P1 and P2) and two battery-powered micropumps. Each *μ*PR was connected to a pressure sensor to measure pressure. P1 was maintained at 1.0 kPa while P2 was varied. We allowed 30 seconds for each P2 stage to provide a sequence of pressures: 1.0 kPa, 1.3 kPa, 1.0kPa, 1.5 kPa, 1.0 kPa, 1.8 kPa, and 1.0 kPa, for a total of 3 minutes and 30 seconds. The liquid-liquid interface between colored streams was recorded with an SMZ-168 stereomicroscope and its camera (Motic Co., ltd., Xiamen, China).

## Results

### A Force-Balance Mechanism Enables a Range of Regulated Outlet Pressure

Pressure regulators are commonly used in pneumatic circuits to reduce high-pressure air to a lower, controllable pressure setpoint for downstream applications. As with most manual pressure regulators, our 3D-printed *μ*PR uses a force-balance mechanism and is designed to maintain a user-defined setpoint suitable for standard microfluidic systems (~1-10kPa). As shown in **Fig. 2**, the *μ*PR consists of a high-pressure air chamber, low-pressure air chamber, and pressure control component. The high-pressure air chamber includes the closing (bottom) cantilever springs, the poppet valve, and the connecting rod. This chamber receives constant pressure from a miniature air pump. The low-pressure chamber with the pressure sensing diaphragm outputs the regulated outlet pressure. The pressure control component consists of 3D-printed top cantilever springs and the control knob (a bolt and a pairing nut), which is used to control the outlet pressure as described below. The operation of *μ*PR can be described in four phases as shown in **Fig. 3**.

**Fig. 2.**
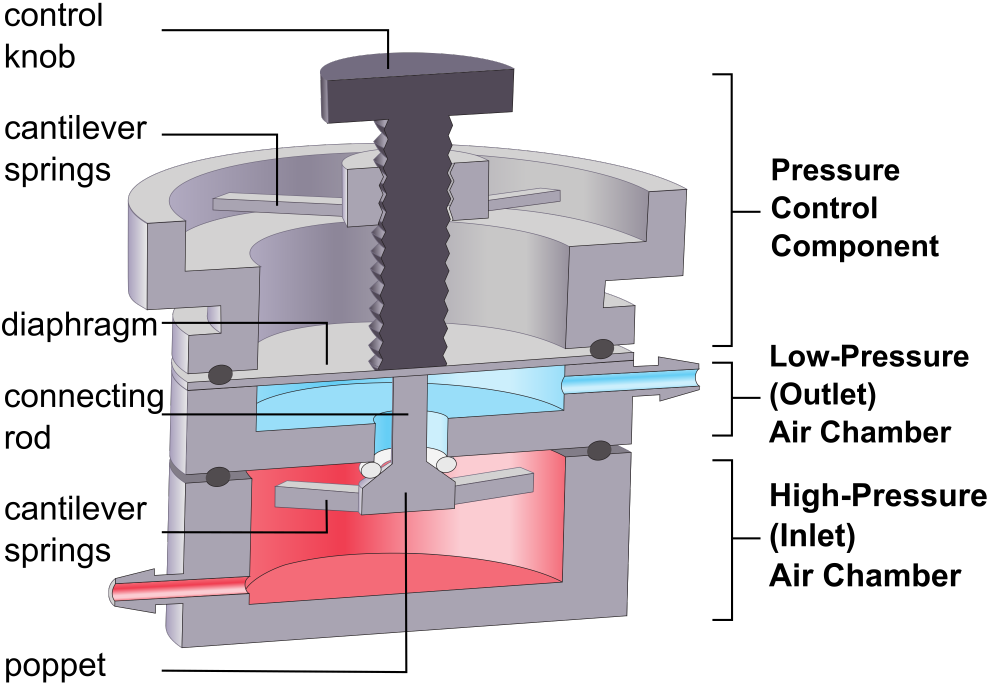
Cross-sectional schematic of essential components in the 3D-printed *μ*PR. The high-pressure chamber (red) receives a constant high-pressure air supply from an external source. The low-pressure chamber (blue) outputs air at a constant low-pressure. The outlet pressure is controlled by adjusting the pressure control component, consisting of cantilever springs and a control knob.

**Fig. 3.**
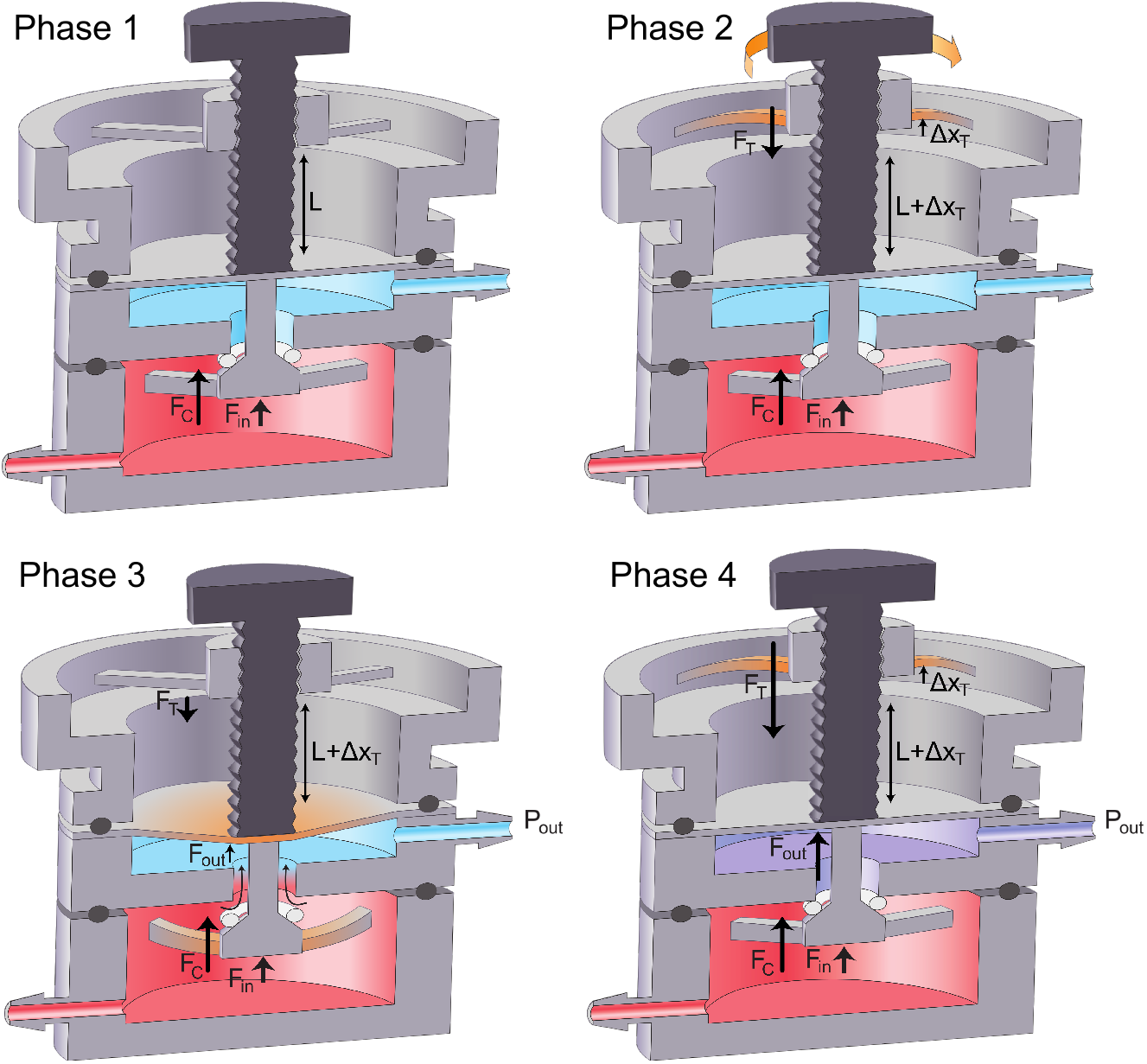
Depiction of the four phases of the pressure regulating process. During Phase 1, the air passage is fully closed, while we supply air at a constant high pressure. In Phase 2, the user turns the control knob to displace the top cantilevers. As the top cantilever restoring force (*F_T_*) increases, the air passage between the chambers remains closed. In Phase 3, when *F_T_* surpasses a certain threshold, the air passage opens. Finally, in Phase 4, the pressure in the low-pressure air chamber will reach the desired level indicated by the control knob and the passage will close. Once the pressure is set by the user, the device toggles between Phase 3 and Phase 4 to maintain the desired output pressure.

#### Phase 1

Constant high-pressure air is supplied to the high-pressure chamber using a miniature air pump. There are two closing forces present at this stage. The inlet pressure force (*F_in_*) is an upward force generated by the inlet pressure acting on the poppet. The closing cantilever spring force (*F_C_*) is a constant upward force generated by the displacement of the non-adjustable bottom cantilever springs and is set during assembly. These upward forces press the poppet to the seat and close the air passage between chambers. In this phase, the bolt length under the nut is *L* and the tip of the bolt rests against the pressure sensing diaphragm without exerting a downward force.

#### Phase 2

As we turn the control knob clockwise, the bolt length under the nut is increased to *L* + Δ*x_T_* and the top cantilever springs are displaced upward from their relaxed state by Δ*x_T_*. This upward displacement of the cantilever springs generates a downward restoring force (*F_T_* = *k_T_* · Δ*x_T_*) on the sensing diaphragm. During this phase, the air passage is still sealed by upward forces (*F_in_* and *F_C_*) because *F_T_* < *F_in_* + *F_C_*.

#### Phase 3

When the control knob is rotated further to increase Δ*x_T_*, *F_T_* overcomes the upward forces (*F_in_* + *F_C_*) and the bolt tip displaces the pressure sensing diaphragm and connecting rod downward. The motion of the connecting rod unseats the poppet valve and opens the air passage, allowing high-pressure air to enter the low-pressure chamber. The pressure (*P_out_*) in the low-pressure chamber exerts an upward force *F_out_* on the bottom surface of the pressure sensing diaphragm (area *A_d_*), *P_out_* = *F_out_* · *A_d_*^−1^.

#### Phase 4

*P_out_* increases until the summation of *F_out_* and other upward forces *F_in_, F_C_* is equal to *F_T_* as shown in **Equation 1**. These upward forces push the poppet valve toward the seat and block air flow between chambers (**Fig. 3**). This allows us to *P_out_* to be set by changing the top cantilever spring force (*F_T_* = *k_T_* · Δ*x_T_*) by adjusting the rotational position of the control knob. Since *P_out_* is used to pressurize a downstream fluid reservoir or channel *P_out_* decreases and the *μ*PR re-enters <u>Phase 3</u> to allow high-pressure air to compensate for the pressure loss. Once Δ*x_T_* is set by the control knob, the *μ*PR toggles between <u>Phases 3 and 4</u> to maintain a stable setpoint, *P_out_*.

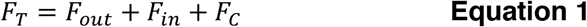

Here, the top cantilever spring force *F_T_* = *k_T_* · Δ*x_T_*; *k_T_* is the spring constant of the top cantilever spring, with Δ*x_T_* being the spring displacement. The outlet pressure force, *F_out_* = *P_out_* · *A_d_*; *P_out_* is the outlet pressure, and *A_d_* is the area of the sensing diaphragm. *F_in_* is the inlet pressure force on the exposed area of the poppet, while *F_C_* is a constant closing force from the bottom cantilever spring.

**Equation 1** simplifies because the closing cantilever springs in the high-pressure chamber are not adjustable, and therefore *F_C_* is a constant. *F_in_* is constant as long as we supply a constant input pressure to the high-pressure chamber. Because both *F_in_* and *F_C_* are constants, we can control *F_out_* (thus *P_out_*) by manipulating the *F_T_* applied to the diaphragm. *F_T_* scales linearly with the displacement (Δ*x_T_*) of the top cantilever springs, hence we can tune *P_out_* by adjusting the angular position of the control knob.

### Calibration of the *μ*PR and Pressure Stability

A major goal of our pumping platform is to provide tunable pressure control while maintaining a portable setup. Therefore, we selected a miniature battery-powered air pump instead of a compressed air line or a pressurized cylinder as the external high-pressure source. Since our *μ*PR operates on the assumption of constant inlet pressure (see **Equation. 1**), we first confirmed that the pressure from the miniature air pump was stable over time. Running at 3 volts, the pump maintained a stable pressure (41 ± 0.02 kPa) over the course of 5 days (see **Fig. S3)**. Next, we sought to characterize the relationship between the angular position of the control knob and the resulting outlet pressure. As shown in **Fig. 4(a)**, we rotated the control knob by 15° increments (corresponding to increasing or decreasing Δ*x_T_* in **Fig. 3**) and measured the output pressure. The data revealed two distinct slopes. In the first region from 1^st^ to the 9^th^ position (1 – 2.1 kPa), the slope was 0.15 kPa per 15° increment while in the second region from the 10^th^ to 20^th^ position (2.3 kPa – 10 kPa) the slope was 0.70 kPa per 15° increment. These different slopes may be a consequence of the compressibility of the sealing O-ring on the poppet valve. That is, the O-ring may be partially in contact with the valve seat and limiting air flow between chambers (positions 1 to 9). With increased rotation (positions 10 to 20), the O-ring detaches fully from the valve seat and air can flow between chambers with less resistance, thus creating a steeper slope relationship.

**Fig. 4.**
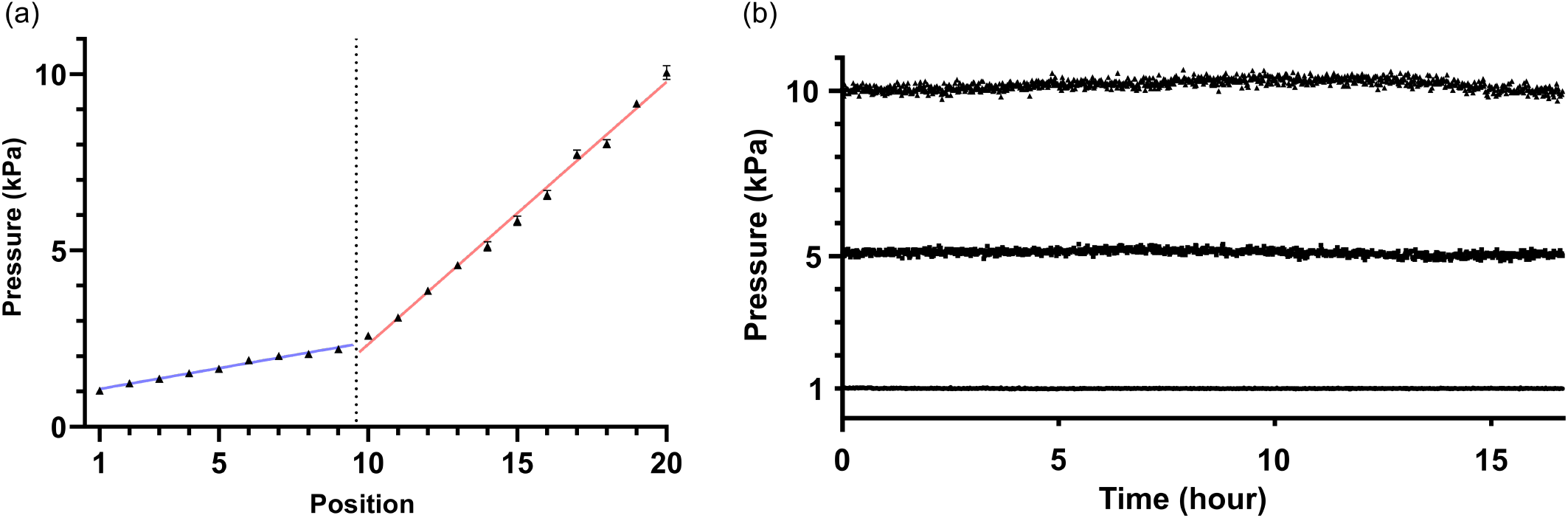
(a) Outlet pressure vs. control knob positions (15° steps). Pressures increase by 0.15-kPa increments between 1^st^ and 9^th^ positions (blue), and 0.70-kPa increments between 10^th^ and 20^th^ positions (red). (b) Outlet pressure stability test with pressures set to 1,5, and 10 kPa, by turning the control knob to the 1^st^, 14^th^, and 20^th^ positions following the calibrated results in (a). The pressure was measured over 16 hours to check the stability of the outlet pressure regulated by the device. The three outlet pressures were 1.0 ± 0.01 kPa, 5.1 ± 0.09 kPa, and 10.2 ± 0.16 kPa throughout the 16-hour test.

To ensure controlled flow for culture applications, it is important to provide a stable pressure drop (Δ*P* = *P_out_* – *P_atm_*) across the microchannel network. Here, the outlet pressure (*P_out_*) regulated by the *μ*PR helps establish Δ*P*. Using the calibration data from **Fig. 4(a)**, we characterized the stability of *P_out_* over 16 hours at three different setpoints, 1, 5, and 10 kPa. As shown in **Fig. 4(b)**, the measured outlet pressures were 1.0 ± 0.01 kPa, 5.1 ± 0.09 kPa, and 10.2 ± 0.16 kPa. The error to measured pressure ratios for 1, 5, and 10 kPa were 1.0%, 1.8%, and 1.6% respectively, demonstrating the *μ*PR’s ability to provide tunable and stable pressures across the output range.

Next, we explored how the *μ*PR could be used to provide a stable pressure drop across a microfluidic channel and produce flow rates practical for cell culture applications. The *μ*PR was designed to support low flow rates that can be difficult to achieve with commercial pressure regulators (e.g., 10 - 100 nL·min^−1^) for cell culture applications. The flow rates were measured in **Fig. 5** for different outlet pressures to investigate the *μ*PR’s capability of controlling the liquid flow. We introduced pressure drops, Δ*P*, from 1 to 8 kPa, using the *μ*PR and measured flow rates ranging from 8.50 nL·min^−1^ to 98.7 nL·min^−1^. We observed an excellent correlation (*R*^2^=0.999) between the COMSOL simulations and experimental flow rate measurements (Δ*P* from 1 to 8 kPa). The slope describing the relationship is 12 nL·min^−1^·kPa^−1^.

**Fig. 5.**
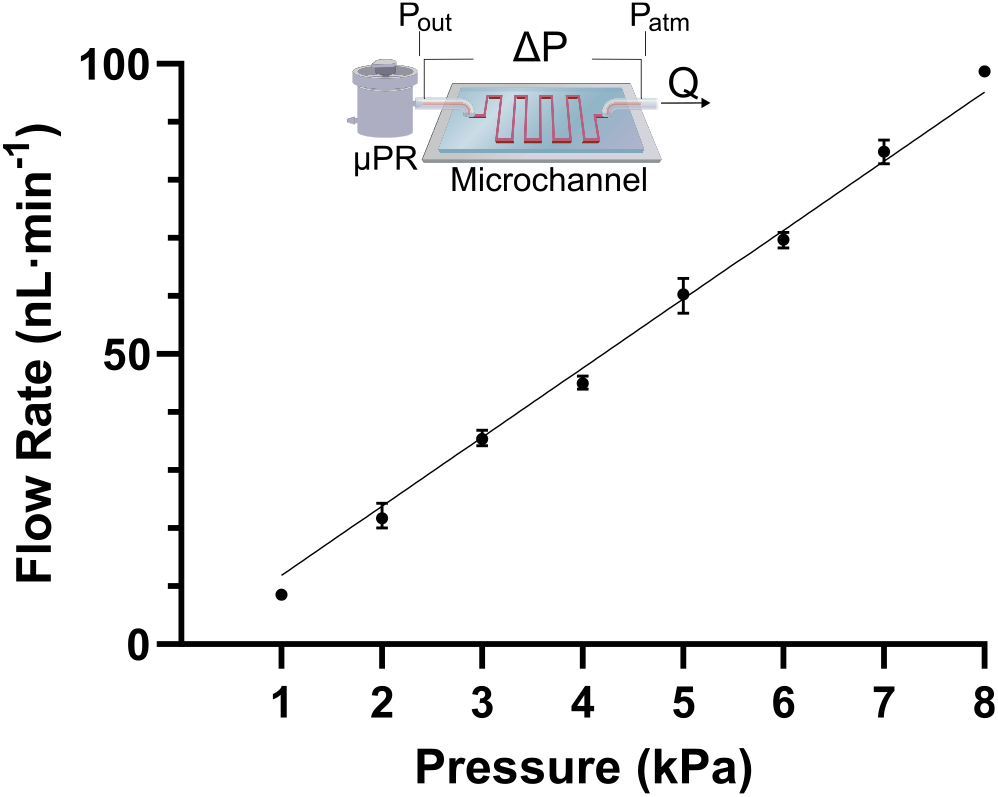
The inset shows the test setup, including the pressure regulator that creates a Δ*P* across the microchannel. Δ*P* is determined by the outlet pressure of *μ*PR and the atmospheric pressure at the end of the microchannel. Δ*P* (1 to 8 kPa) covers flow rates from 9 to 100 nL·min^−1^, yielding a slope of 12 nL·min^−1^·kPa^−1^. The straight line is drawn with the simulated response of flow rates vs. the outlet gauge pressures. *R*^2^ = 0.999 is the correlation between the experimental data and the COMSOL simulation results.

### Microfluidic Culture of Human Umbilical Vein Endothelial Cells (HUVECs)

In microfluidic systems, media perfusion is required because the small media volume in the channel is rapidly depleted of nutrients by metabolically active cells and must be replenished to maintain cell viability. To demonstrate the compatibility of our *μ*PR to control fluid flow and maintain cells, we used the *μ*PR to establish an endothelial monolayer in a tissue barrier model that we previously developed (Mansouri et al. 2022). As shown in **Fig. 6a**, the culture platform consists of two microchannels separated by a nanomembrane. The lower channel was filled with cell media while the top channel was supplied with flows driven by the *μ*PR. The *μ*PR induced a stable pressure drop of 8 kPa across the top culture microchannel, resulting in a constant 1*μ*L·min^−1^ flow rate for introducing cell media from the reservoir into the culture region.

**Fig. 6.**
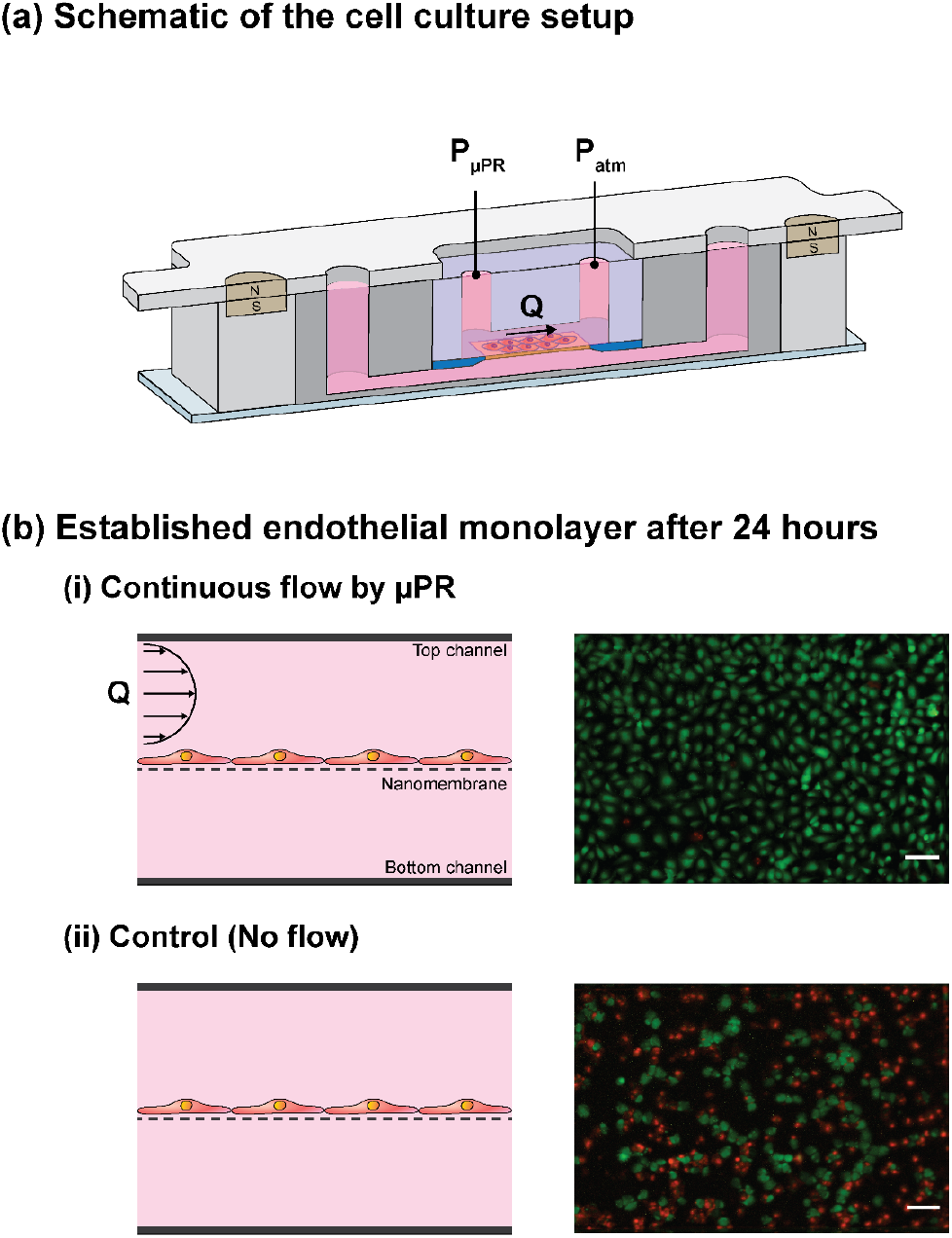
(a) Schematic illustration of the cell culture platform. A mini air pump supplies high-pressure air to *μ*PR, which outputs a stable pressure drop (Δ*P*) across the top microchannel of the platform. This results in the flow of cell media from the reservoir into the microchannel. The platform consists of two microchannels separated by an ultrathin nanomembrane. Components of the platform can be disassembled after the experiment due to its reversible magnetic latching mechanism. We set the output of 8 kPa from the *μ*PR to drive the culture media flow (Q = 1 *μ*L·min^−1^). (b) Cross-sectional view of the endothelial monolayer, and comparison of cultured cells in (i) dynamic culture (with the flow) and (ii) static culture (no flow). The cells were stained with LIVE/DEAD stain and fluorescence images were captured in green (viable cells) and red (dead cells). This demonstrates that the *μ*PR can drive continuous flow vital for long-term cell culture and the formation of a confluent cell monolayer. Scale bars = 100*μ*m.

As expected, cells cultured in the device with media flow driven by the *μ*PR were maintained alive and formed a confluent monolayer after 24 hours while the majority of cells in the static control died due to lack of cell media supply (**Fig. 6b**). The live/dead staining showed a 98.5% survival rate in the *μ*PR-supplied device whereas the static control (no media flow) had a 38.2% survival rate. These results confirmed the capability of the *μ*PR to deliver stable flow rates and maintain a long-term culture of cells in microfluidic devices.

### Dynamic pressure control using the control knob

Since the outlet pressure can be easily changed based on the calibrated position of the control knob, we demonstrate *μ*PR’s responsiveness to real-time pressure switching. Here we show responses that that changes can be made to the ranges of pressure – low to high, medium to high, and low to medium. As shown in **Fig. 7**, we measured dynamic pressure adjustments, each with three stages, that spanned the tested range: i) 1 kPa - 5 kPa - 1 kPa, ii) 5 kPa - 10 kPa - 5 kPa, and iii) 1 kPa - 10 kPa - 1 kPa. In this experiment, we again used the calibration results as presented in **Fig. 4(a)** for setpoints of the control knob positions for pressures used in this experiment. **Fig. 7** shows that our *μ*PR could ramp up and down to reach desired setpoints within one-minute periods, even among the largest dynamic pressure patterns in the experiment.

**Fig. 7.**
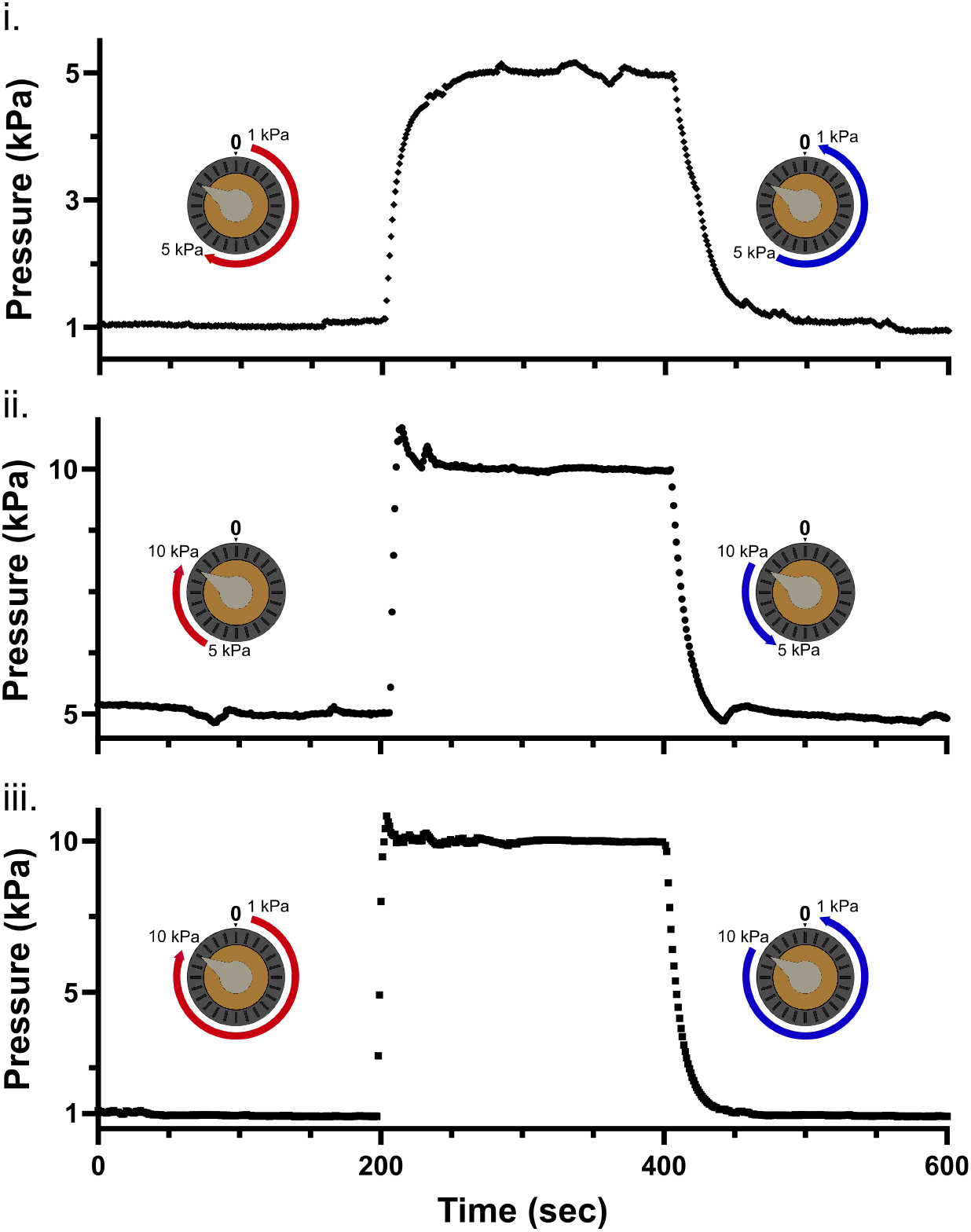
Dynamic responses of pressure patterns including, (i). 1 to 5 to 1 kPa, (ii). 5 to 10 to 5 kPa, and (iii). 1 to 10 to 1 kPa are achieved via control knob turns with the calibration data in **Fig. 4(a)**. Each pattern features three stages, each with 200 seconds under the real-time observation of the dynamic pressure response.

To highlight the integration of multiple *μ*PRs in a single system, we utilized two *μ*PRs to separately control the flow rates of two liquids within a Y-shaped microfluidic channel and visualized the dynamic equilibrium position of the dual-stream laminar flow interface while adjusting one *μ*PR to a new setpoint. We fed red-dyed deionized water to the top inlet port of the Y-channel with the pressure set to 1.0 kPa *μ*PR, P1. Blue-dyed deionized water was fed into the bottom inlet port with pressure regulated by a second *μ*PR, P2; these pressure values were changed during the experiment from a range of 1.0 kPa to 1.8 kPa.

As expected, when P1 = P2, the liquid-liquid interface between the red and blue streams was located at the midline of the channel (white dashed line), indicating *μ*PRs’ capability of delivering stable flow rates using multiple *μ*PRs. As we changed P2 from 1.0 kPa to 1.8 kPa by turning the control knob, the flow rate in the bottom channel increased and the interface was shifted upward (see **Fig. 8** and **VIDEO S4)**. We allowed a 30-second period of observation time for each new P2 set point with the following sequence of pressures: 1.0 kPa, 1.3 kPa, 1.0kPa, 1.5 kPa, 1.0 kPa, 1.8 kPa, and 1.0 kPa, for a total of 3 minutes and 30 seconds. The liquid-liquid interface shifted in response to the P2 pressure adjustment and quickly settling to the new position, and maintained stability during each of the 30-second pressure monitoring periods. The dynamic response of the *μ*PR flow adjustment demonstrated real-time pressure adjustment and stable dynamic equilibrium positions. We highlighted the pressure control capabilities of the system and flows profile possibilities for more advanced real-time features that require pressure controls.

**Fig. 8.**
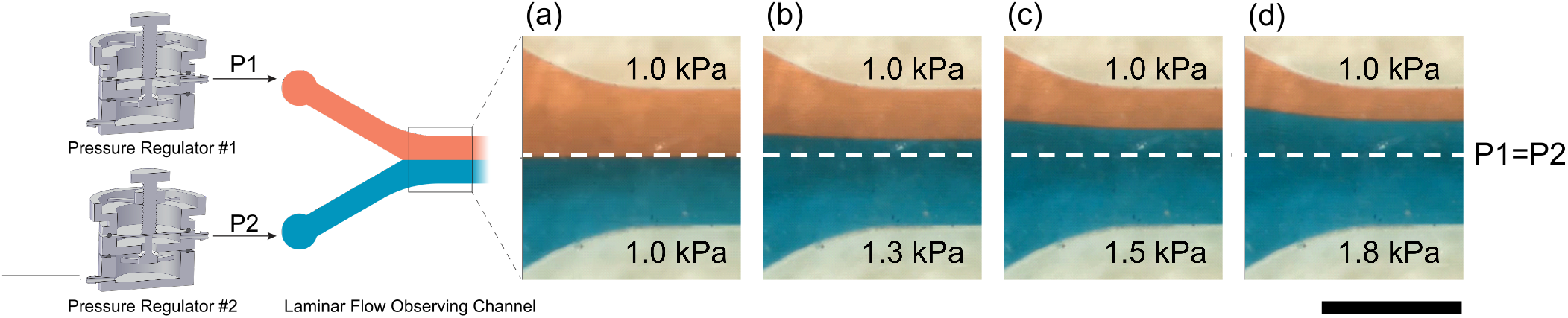
Real-time observation of co-laminar liquid flow pressurized with two *μ*PRs. *μ*PR #1 supplies pressure (P1) to one inlet port of the laminar flow observing channel, while *μ*PR #2 supplies pressure (P2) to the other. P1 is set to 1 kPa as control, while P2 is adjusted to (a) 1.0 kPa, (b) 1.3 kPa, (c) 1.5 kPa, and (d) 1.8 kPa using the control knob. Scale bar = 1 mm

## Discussion

The goal of our platform is to provide a portable, simplified microfluidic flow control method while providing stabile flows suitable for cell culture applications. While there are commercial solutions for pneumatic pressure control, these pressure regulators have larger footprints (>30mm), a higher outlet pressure range (~35 kPa) with a lower resolution (>3.5 kPa resolution), cannot be customized, are expensive (>$100 USD for one with aforementioned features), and require a laboratory compressed air line. By introducing the *μ*PR along with a mini air pump to create a microfluidic flow control platform, we can deliver a range of tunable and stable flow rates within a portable system size. Our platform provides a cost-effective pressure control scheme with a range of customization opportunities owing to the increasing availability of hobby and commercial 3D printers. For reference, the total cost of the mini air pump and *μ*PR setup as shown in this work is less than $7 USD, of which the *μ*PR is less than $1.20 as shown in **Table S5**.

In our design, (see **Fig. 2** and **Fig. 3)**, the pressure regulating mechanism is similar to that of conventional pressure regulators. However, by incorporating 3D-printing techniques, we were able to integrate two sets of cantilever springs as an alternative to large commercial springs to simplify the assembly and help miniaturize the device. By incorporating cantilever springs into the poppet valve design, we created an upward closing force (*F_C_*), as shown in **Fig. 3**, to prevent possible high-pressure air leakage to the low-pressure chamber through the air passage. This “normally-closed” design allows users to shut off output pressure and momentarily disconnect the cell culture compartments for inspection or modification. Since regulation of *P_out_* depends on the closing actions of the poppet valve, we chose a gas-impermeable elastomeric Viton O-ring (shore 60A) at the poppet for better sealing. This suits our target applications, which are often operated with a low-pressure and flow rate regime. To target the range of 1-10 kPa, we chose the M2 size (0.4-mm pitch, 2-mm diameter) bolt as the control knob and partner it with a 24-position dial. Such a combination provides sufficient pressure resolution (< 1kPa per 15° turn) while retaining user-friendly control. By adjusting some key mechanical parameters, such as *k_T_* and *A_d_*, we can achieve different targeted outlet pressure ranges. **Equation 2** shows that *k_T_* can be modified by changing the mechanical properties of the material by either switching to a different material or changing the curing settings of the 3D printer. *k_T_* can also be altered by the geometry of cantilevers. For instance, we can adjust the pressure response by decreasing *k_T_* by increasing the length of the cantilever springs or decreasing their width or thickness, as shown in **Equation 2**.

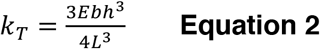

Where *E* is the Young’s modulus of the 3D-printed material, and *b, h, L* are the width, thickness, and length of each cantilever, respectively.

Although *k_T_* is more sensitive to changes in the thickness (*h*) of the cantilever than the width (*b*) (see **Equation 2**), the z accuracy (i.e., layer thickness control) of the 3D printer is often less than the x-y axes, resulting in greater variability in the thickness (Melenka et al. 2015). The 3D-printed *μ*PR can be modified to fit different pressure ranges. For example, increasing the diameter of the sensing diaphragm *A_d_* can improve the resolution of the output pressure setpoint but results in a larger device footprint and a smaller upper bound (constrained by maximum cantilever spring force) of the outlet pressure, since the outlet force scales linearly with the diaphragm area (*F_out_* = *P_out_* · *A_d_*) but is limited by the top cantilever spring force.

3D-printed structures are still associated with dimensional errors for such small device features, thus the calibration of the *μ*PR’s outlet pressure is device-specific. The relationship between control knob positions and outlet pressures, once calibrated, can be used to produce the desired outlet pressure. The *μ*PR does not come into contact with the fluid and can be reused as needed. The compact and easy setup of the *μ*PR-based microfluidic flow control platform provides manual control of Δ*P* based on the calibration. Using this culture platform, we were able to deliver a constant flow rate of media to a culture platform to maintain a viable environment for HUVECs as compared to the no-flow situation. With the dynamic control capability demonstrated with the co-laminar flows, we present more possibilities in dynamically controlling the outlet pressure to introduce different media flow rates for culture setup changes using our *μ*PR (e.g. shear stress adjustment for cell alignment purposes) without modifying the channel geometry. In contrast to syringe pumps and commercial pneumatic solutions, the small footprint and minimal peripheral equipment requirements of the *μ*PR-based system and be easily moved in and out of a cell culture incubator.

## Conclusions

In summary, we introduced an easy-to-fabricate, low-cost miniaturized 3D-printed pressure regulator and highlighted stable pressure control capabilities for microfluidic applications. We anticipate that our open access design files and simple fabrication techniques will enable other laboratories to customize *μ*PR designs to support a broad range of microfluidic applications where syringe pumps or traditional pneumatic methods are not appropriate.

## Supporting information

laminar flow control video (S4)

## Acknowledgments

This work was supported in part by the National Institute of Health under award numbers 1R43GM137651 and 1R61HL154249. The authors thank Xian Boles from the RIT Medical Illustration MFA program for illustration support.

