## Supplemental figures for "A Miniaturized 3D-Printed Pressure Regulator (*μ*PR) for Microfluidic Cell Culture Applications"

Supplementary:

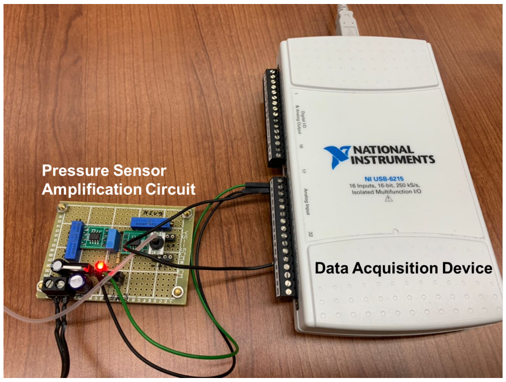


**Fig. S1.** We utilized a pressure sensor, (TBPDANS005PGUCV, Honeywell International Inc., Charlotte, NC, USA), a signal amplification circuit, and a data acquisition device, (NI USB-6215, NI, Austin, Texas, USA) to record the outlet pressure of the device. The pressure reading unit was calibrated using a commercial pressure regulator (Type-90, ControlAir, Inc., Amherst, New Hampshire, USA) and a pressure gauge (DPG-110, Dwyer Instruments, Inc., Michigan City, Indiana, USA).

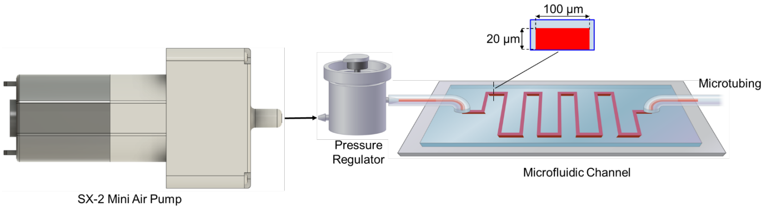


**Fig. S2.** The driven flow rate test setup consists of the SX-2 mini air pump, the 3D-printed pressure regulator, the microfluidic channel, and a microtubing that collects driven mass of fluids to calculate the flow rates with travel times.

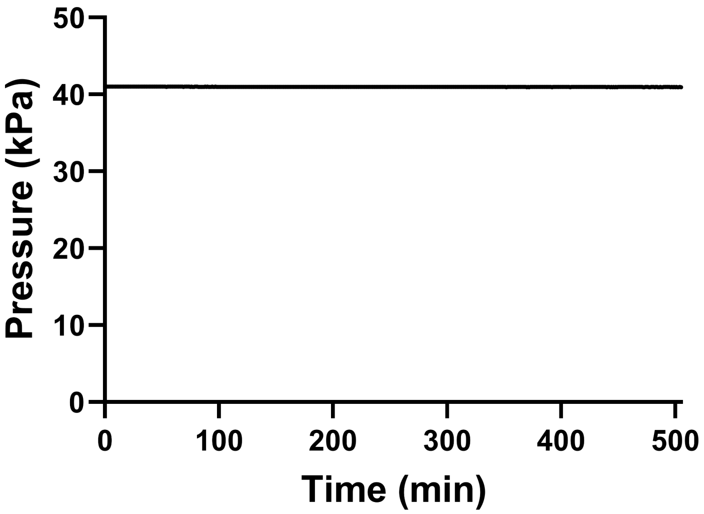


**Fig. S3.** We recorded the pressures supplied by mini air pump powered at 1.5V, 0.14A over 500 minutes to verify the mini air pump's capability of supplying stable pressure as the inlet for the pressure regulator. The results demonstrated a stable 41-kPa outlet pressure over the examined period.
